# Peptidoglycan remodeling in response to cell wall acting antibiotics in *Bacillus subtilis*

**DOI:** 10.1101/2023.01.23.525174

**Authors:** Charlène Cornilleau, Laura Alvarez, Christine Wegler, Cyrille Billaudeau, Felipe Cava, Rut Carballido-López

## Abstract

Most bacteria are encased into a load-bearing rigid framework, the cell wall (CW). The peptidoglycan (PG) layer, a network composed of glycan strands cross-linked by stem peptides, is the main component of the CW. During PG synthesis, precursors are first synthetized intracellularly, before being incorporated into the existing PG meshwork by transglycosylation (TG) and transpeptidation (TP) reactions. Covalent modifications of the PG meshwork such as amidation and acetylation participate in PG homeostasis by regulating PG-associated enzymes like PG hydrolases.

Because of its essential role, PG synthesis represents a primary target for antibiotic action. Here, we investigated the effect on PG composition of antibiotics targeting intracellular and extracellular steps of PG synthesis: inhibitors of PG precursors synthesis (fosfomycin, D-cycloserine, bacitracin and tunicamycin) and TG/TP inhibitors (vancomycin and penicillin G), respectively. Our study revealed interesting correlations between crosslinking and both de-N-acetylation and amidation of the sacculus. A thorough analysis of muropeptides composition put into light an unexpected anti-correlation between the degree of PG crosslinking and accumulation of de-amidated disaccharide-tripeptide monomer subunit (M3) in the presence of TP inhibitors. We confirmed these observations by analyzing mutants of the PG synthesis pathway.

## Introduction

The bacterial cell wall (CW) is a complex, semi-rigid extracellular meshwork that maintains shape and provides structural integrity to bacterial cells. The principal constituent of the CW is peptidoglycan (PG, also known as the murein sacculus): a three-dimensional network of linear sugar strands of alternating N-acetyl-glucosamine (GlcNAc) and N-acetyl-muramic acid (MurNAc) cross-linked by stem peptides attached to MurNAc. The stem peptides are synthesized as pentapeptide chains and contain the unusual D-amino acids. In most Gram-positive species the pentapeptide chain is typically made of L-alanine (L-Ala) in position 1, D-glutamine or D-iso-glutamic acid (D-Glu) in position 2, L-lysine in position 3 and a C-terminal D-alanine-D-alanine (D-Ala-D-Ala) dipeptide (Strange et Kent 1959; Mandelstam et Rogers 1959; Vollmer, Blanot, et De Pedro 2008; Pazos et Peters 2019). However, in Bacilli as well as in Mycobacteria and Gram-negative species, the lysine in position 3 is substituted by *meso*-2,6-diaminopimelic acid (*m*-DAP), leading to the following stem peptide sequence for the Gram-positive model bacterium *Bacillus subtilis*: L-Ala, D-Glu, *m*-DAP, D-Ala, D-Ala.

PG synthesis results from a series of sequential reactions that occur in three overall stages. First, the soluble, activated nucleotide PG precursors UDP-GlcNAc and UDP-MurNAc*-*pentapeptide are synthesized in the cytoplasm by the sequential action of the Mur enzymes. Then, the soluble precursors are transferred to undecaprenyl phosphate (UP, also referred as C_55_ lipid carrier) at the inner leaflet of the membrane to finally form the lipid-anchored GlcNAc-MurNAc-pentapeptide monomer subunit, namely lipid II, which is then flipped across the membrane. Last, externalised lipid II is polymerised into a nascent glycan chain by transglycosylation (TG) and, concomitantly, crosslinked to the existing sacculus by transpeptidation (TP). **Figure 1** displays the PG synthetic pathway and known enzymes in *B. subtilis*. Historically, most of the enzymes responsible for TG and TP reactions were discovered because of their ability to bind penicillin, and were thus called penicillin-binding proteins (PBPs). PBPs are classified depending on their activities: Class A and B high-molecular weight PBPs are PG synthases, and Class C low molecular-weight PBPs are PG hydrolases. Class A PBPs (aPBPs) possess two functional domains and perform both TG and TP reactions, while Class B PBPs (bPBPs) are monofunctional and only have a TP catalytic domain. In addition to aPBPs, the penicillin-insensitive essential RodA and FtsW proteins also catalyze TG from lipid II, working with their cognate bPBPs TPs to synthesize sidewall and septal PG, respectively (Emami et al. 2017; Taguchi et al. 2019). TP reactions by aPBPs and bPBPs cleave the terminal D-Ala from a pentapeptide and form a peptide bridge between the D-Ala in position 4 of the acyl-donor peptide and the D-center of the *m*-DAP in position 3 of the acyl-acceptor stem peptide (**Fig. 1**). This 4→3 crosslink, termed D,D-transpeptidation (DD-TP), strictly requires a pentapeptide as substrate (Terrak et al. 1999) and is the main crosslinking strategy in most bacteria including *B. subtilis* (Warth et Strominger 1971; Atrih et al. 1999; Vollmer, Blanot, et De Pedro 2008).

**Figure 1.**
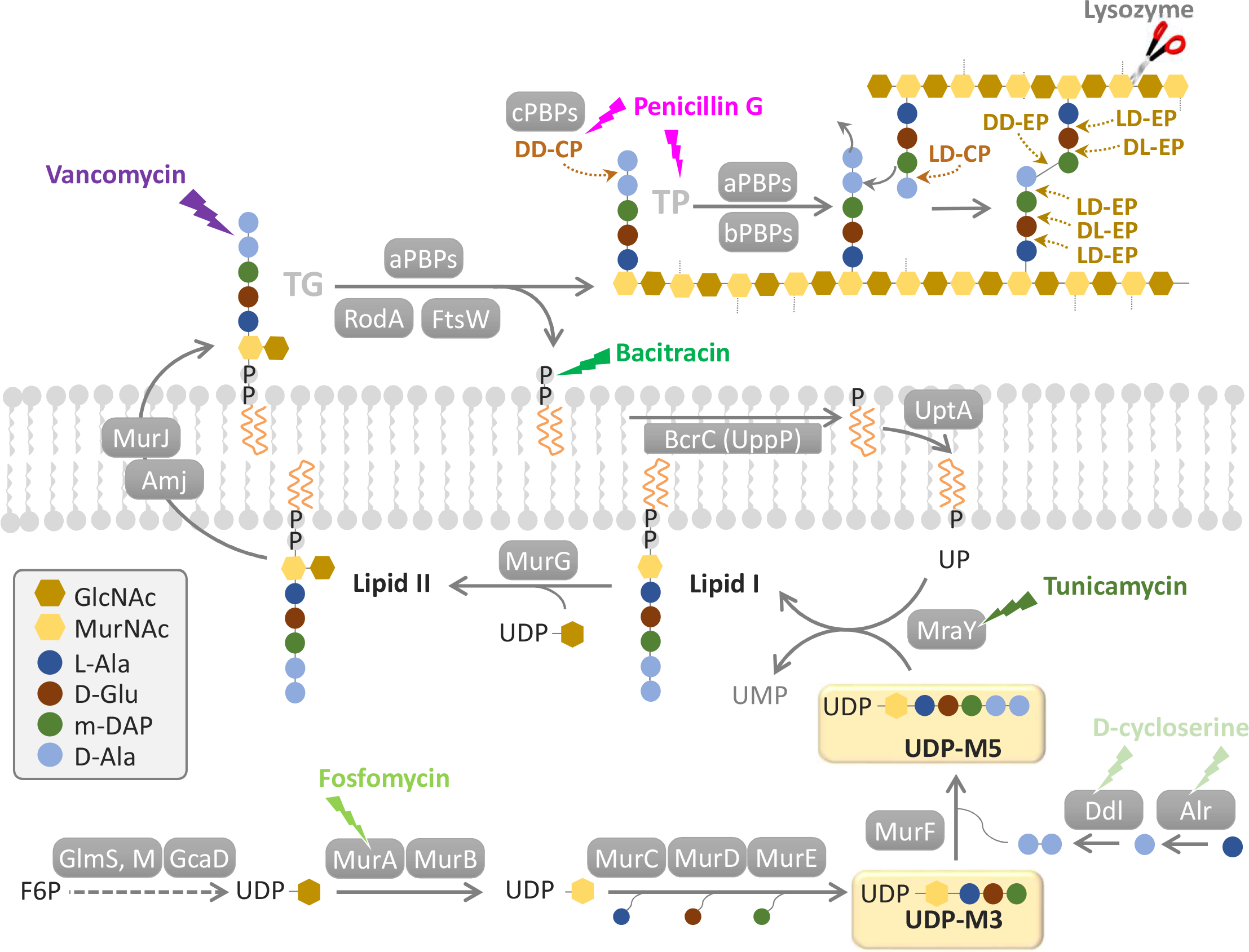
Schematic view of the PG synthesis pathway in *B. subtilis*. The first step is the synthesis of the nucleotide-activated UDP-GlcNAc from fructose-6-phospate (F6P), which is then transformed into UDP-MurNAc by the MurA and MurB enzymes. UDP-MurNAc-tripeptide (UDP-M3) is next formed by sequential addition of L-Ala by MurC, D-Glu by MurD and *m*-DAP by MurE. The D-Ala-D-Ala dipeptide, synthesized by the alanine racemase Alr and the D-Ala-D-Ala ligase Ddl is then added by MurF to produce UDP-MurNAc-pentapeptide (UDP-M5). MraY catalyzes then the first committed membrane-bound step by transferring UDP-M5 onto UP to form the lipid I intermediate at the inner leaflet of the cytoplasmic membrane. Then, MurG adds a GlcNAc unit to lipid I to form lipid II, which is then flipped to the outer leaflet of the cytoplasmic membrane by the major (MurJ) and the minor (Amj) flippases (Meeske et al. 2015). The externalised PG precursor is linked to a nascent glycan strand by enzymes possessing transglycosylase activity, releasing undecaprenyl-pyrophosphate (UPP) in the process. The released UPP will be dephosphorylated to UP by UPP phosphatases (BcrC, UppP, and PgpB (YodM) to a minor degree, (Radeck, Lautenschläger, et Mascher 2017)) and flipped back to the inner leaflet of the membrane by the recently identified UptA (Roney et Rudner 2022), allowing the recycling of the lipid carrier for another round of PG precursor transport. Finally, the nascent glycan chain is incorporated to the existing sacculus by TP reactions between stem peptides (Sauvage et al. 2008). PG hydrolases are needed to open the network and allow growth: D,D-carboxypeptidases (DD-CP), L,D-carboxypeptidases (LD-CP), D,D-endopeptidases (DD-EP), D,L endopeptidases (DL-EP). PG synthesis is the target of antibiotics inhibiting the synthesis of PG precursors (fosfomycin, D-cycloserine, tunicamycin, bacitracin) and TG/TP reactions (vancomycin, penicillin G).

In addition to PG synthases, growth of the sacculus requires the activity of PG hydrolases, which cut existing bonds to allow the insertion of new material and thus the expansion of the PG meshwork without increasing its thickness (Typas et al. 2012). PG hydrolases can cleave almost any glycoside and peptide bond and belong to three different families depending on their substrate: (i) glucosaminidases, muraminidases, lytic transglycosylases, and also the lysozymes produced by many organisms as a major effector of their innate immune system, cleave the sugar backbone, (ii) amidases cut the bond between MurNAc and the stem peptide, and (iii) peptidases hydrolyze peptide bonds, either within or between peptide stems (Vollmer et al. 2008; Hashimoto et al. 2018). Peptidases include Class C low molecular-weight PBPs (cPBPs) and can be further classified as carboxypeptidases (CPs) or endopeptidases (EPs) depending on whether they remove the carboxy-terminal amino acid of the stem peptide or cleave non-terminal peptide bonds, respectively. Peptidases can also be classified depending on their stereospecificity, whether they cleave the bond between two D-chiral centers (DD-EPs and DD-CPs) or between and L- and a D-chiral center (LD- and DL-EPs and CPs) (**Fig. 1**). PG crosslinking and the potential lethal activity of PG hydrolases (autolysis) are also regulated by covalent modifications of their substrate (either the glycan strands or the stem peptides). Common PG modifications known to modulate PG hydrolases include O-acetylation of MurNAc residues (Guariglia-Oropeza et Helmann 2011; Yadav, Espaillat, et Cava 2018) and also of GlcNAc in some species (Moynihan et Clarke 2011; Bernard et al. 2012), de-N-acetylation of GlcNAc (Atrih et al. 1999; Yadav, Espaillat, et Cava 2018) and de-amidation of D-Glu or *m*-DAP (Vollmer, Blanot, et De Pedro 2008; Dajkovic et al. 2017). The detailed composition of the vegetative sacculus of *B. subtilis* was reported just before the end of the last century and emphasized the de-amidation of *m*-DAP and the de-N-acetylation of GlcNAc as main PG modifications, and showed a role in PG maturation for the major DD-CP PBP5 (encoded by *dacA*) (Atrih et al. 1999).

PG modifications are part of a complex regulatory response and underlay the plasticity of the PG sacculus to adapt to a changing environment and/or to face conditions challenging its structural integrity. Among the latter, antibiotics targeting the CW are on the front line. Because of its essentiality and also because it is specific to bacteria, the bacterial CW and in particular PG synthesis represent the most prominent target of antibiotics. Almost every step of PG synthesis is targeted by at least one known antibiotic (Schneider et Sahl 2010) (**Fig. 1**). For example, fosfomycin blocks the first committed cytoplasmic step of PG synthesis by covalently binding to the active site of MurA, which inactivates the enzyme (Aghamali et al. 2019; Dörries, Schlueter, et Lalk 2014; Kahan et al. 1974). D-cycloserine is a structural analogue of D-Ala and targets different enzymes that use D-Ala as substrate (Neuhaus et Lynch 1964; Lambert et Neuhaus 1972). It inhibits Alr and Ddl, synthesizing the D-Ala-D-Ala dipeptide (Walsh 1989; Batson et al. 2017). Recently, it was proposed that D-cycloserine can also inhibit transpeptidases (Kuru et al. 2019). Tunicamycin blocks PG synthesis by competing with UDP-M5 to bind MraY (Liu et Breukink 2016; Noda et al. 1992), and it has also been shown to inhibit TG/TP reactions (Ramachandran et al. 2006). Bacitracin is an antimicrobial peptide that binds to the UPP released upon PG precursor TG, preventing its dephosphorylation and thus inhibiting the recycling of the lipid carrier (Radeck et al. 2016; Stone et Strominger 1971). Vancomycin is also an antimicrobial peptide that binds the terminal D-Ala-D-Ala dipeptide of the externalised PG precursors, preventing TG and TP by steric hindrance (Mainardi et al. 2008). Finally, the beta-lactam ring of penicillin G (and beta-lactam compounds in general) mimics the D-ala-D-ala dipeptide, inhibiting PG crosslinking by blocking DD-TPs and DD-CPs (Schneider et Sahl 2010).

Although the primary targets of CW antibiotics are known, the events resulting from primary target inhibition and leading to either adaptation or cell death remain largely unknown. Among these, changes in the chemical composition and structure of the sacculus induced by CW antibiotics might reflect mechanism modulating CW properties. Given the rising emergence of multi-resistant bacteria, it is imperative to comprehensively study the effect of antibiotics in model organisms and use this knowledge for the development of new tools to combat pathogenic bacteria. Here, we have performed the first comprehensive analysis of the effect of PG-targeting antibiotics in the composition and structure of the bacterial CW. Using a high-throughput approach for the preparation and analysis of PG samples (Hernandez et al. 2022), we studied the changes in the sacculus of *B. subtilis* in response to exposure to lysozyme and to six antibiotics targeting different steps of the PG biosynthetic pathway at different sub-lethal concentrations and growing conditions. Notably, our study reveals that cells adapt to the presence of PG synthesis inhibitors: after long exposure times, cultures reach similar optical densities as untreated cultures and restore their pool of intracellular soluble PG precursors. However, the amount of PG in the walls of antibiotic-treated cells is strongly reduced and its structure significantly altered: sacculi are less cross-linked, more N-acetylated and less amidated, suggesting that the adaptative response involves activation of PG hydrolases. We also found interesting correlations between crosslinking and both de-N-acetylation and amidation of dimers. Remarkably, we additionally observed that the dose-dependent decrease in crosslink produced by TP inhibitors was correlated with the accumulation of de-amidated disaccharide-tripeptide monomer subunit (M3). Analysis of individual mutants affected in different aspects of PG metabolism provided some insight into the de-N-acetylation mechanism and uncovered a novel pathway for muropeptide de-amidation. Importantly, our study highlights a key role for endopeptidases in CW antibiotics tolerance and provides additional insights into the mechanisms of PG homeostasis.

## Results and discussion

### Experimental design

To gain insight into the mechanisms regulating PG homeostasis, we aimed at analysing the muropeptide profile of *B. subtilis* in response to a comprehensive set of antimicrobials targeting different steps of PG synthesis (**Fig. 1**). The synthesis of soluble PG precursors was inhibited by fosfomycin and D-cycloserine. Synthesis of the lipid-linked PG precursors was inhibited by tunicamycin and bacitracin. Note that tunicamycin primarily blocks WTAs synthesis but at the high concentrations used here (> 10 μg/ml) it additionally blocks PG synthesis (Brandish et al. 1996; Campbell et al. 2011). Polymerisation and crosslinking of the PG meshwork was targeted by vancomycin and penicillin G. We also analysed the effect of the hydrolytic enzyme lysozyme, and used the antibiotic chloramphenicol, which blocks protein synthesis by targeting ribosomes but does not impact cell surface expansion (Kitahara et al. 2021), as negative control. We aimed to investigate the direct responses as well as the adaptation of *B. subtilis* to CW targeting antibiotics. To this end, cells were treated with sub-lethal concentrations of the compounds either at early-exponential growth (OD_600_ ≈ 0.2) during 2h (‘2h protocol’) or at very low density (OD_600_ ≈ 0.01) during 17h (‘overnight protocol’) (**Fig. S1A**) (See M&M for the detailed procedures). Seven increasing sublethal concentrations of each compound were used for each protocol for muropeptide analysis (**Fig. S1, Fig. S2** and **Fig. S3**). We also extracted and quantified the soluble PG precursors from cultures treated with one low and one high antibiotic concentration (**Fig. S1**).

### Cells adapt to grow in the presence CW antibiotics with little amount of PG

Addition of PG synthesis inhibitors to actively growing cells had a quick impact in growth rate. Upon addition of the compounds, cultures stopped growing for 15-30 min and then often lysed for 30-120 min before resuming growth again for all compounds excepting tunicamycin (**Fig. 2A, Fig. S2C-J** top panels). The concentrations of tunicamycin used to inhibit PG synthesis in this study strongly inhibit WTAs synthesis as well, which may explain the different behaviour. For all other compounds, an adaption to grow with an impaired PG biosynthetic pathway was ultimately implemented and cells started growing again. Viability and/or growth defects of several mutants affected in different aspects of CW synthesis can be rescued by millimolar concentrations of magnesium in the growth medium (Formstone et Errington 2005; Chastanet et Carballido-Lopez 2012), possibly through an inhibition of autolysins by Mg^++^ (Dajkovic et al. 2017; Tesson et al. 2022). When we supplemented the cultures with 20 mM MgSO_4_, lysis was inhibited and/or growth rate improved (**Fig. S2**; bottom panels), suggestive of unbalanced PG hydrolytic activity in the presence of the PG inhibitors. For penicillin G-, vancomycin- and tunicamycin-treated cells, the compensatory effect of Mg^++^ was very mild and only effective at the lowest concentrations of the antibiotics (**Fig. S2F-H**), indicative of a stronger or different deregulation of PG hydrolases in these conditions. After 2h treatment with CW antibiotics, the amount of PG per cell ranged between 17% and 80% of the untreated control, and only the chloramphenicol treated control cells contained normal amounts of PG (**Fig. 2B, Fig. S4**). For antibiotics inhibiting the synthesis of the intracellular soluble precursors, the concentration of UDP-M3 and UDP-M5 species was consistent with the antibiotic target (**Fig. 2C** and **Table S1A**). In fosfomycin-treated cells, UDP-M3 was strongly depleted. Bacitracin, by inhibiting the recycling of lipid carrier, induced the accumulation of UDP-M5 (1.7 to 6-fold). Meanwhile, D-cycloserine - by inhibiting the formation of the D-Ala-D-Ala dipeptide-induced an impressive accumulation of UDP-M3, up to 320-fold for the high concentration. Inhibition of lipid I synthesis by tunicamycin brought about the expected accumulation of UDP-M5 and to a lesser extent of UDP-M3. As previously reported (Vemula et al. 2017), antibiotics targeting TG and TP reactions also affected the concentration of intracellular soluble muropeptides. Vancomycin produced a massive accumulation (118-fold) of UDP-M5 (Cegelski et al. 2002; Singh et al. 2017)and also an important accumulation of UDP-M3 (13-fold) (**Fig. 2C** and **Table S1A**). By blocking newly externalised lipid II, vancomycin may prevent recycling of the lipid carrier to which UDP-M5 should be transferred to form new lipid I and II, leading to accumulation of UDP-M5 in the cytoplasm, and thus of the upstream UDP-M3 too. Penicillin G also induced an accumulation of both UDP-M5 and UDP-M3 (**Fig. 2C** and **Table S1A**). Interestingly, antibiotic-induced accumulation of the intracellular soluble UDP-M5 and/or UDP-M3 precursors affected mostly their non-amidated forms (**Table S1**). Since all lipid II is externalised in its amidated form (Atrih et al. 1999), this is consistent with the intracellular membrane-bound PG precursors being the preferential substrate for *m*-DAP amidation (Levefaudes et al. 2015; Vollmer, Blanot, et De Pedro 2008).

**Figure 2.**
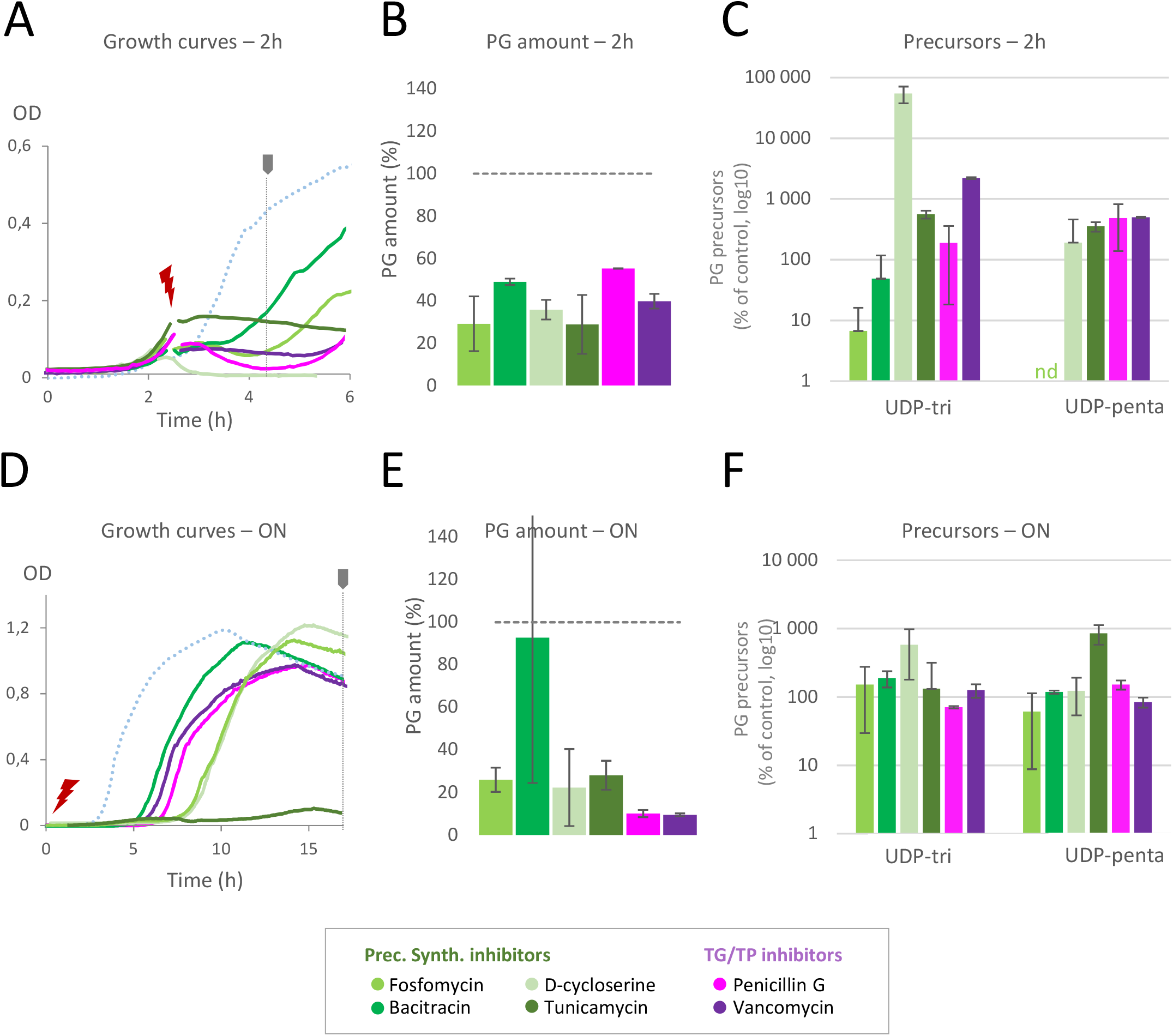
Quantification of total PG and soluble PG precursors upon antibiotic treatment. Results shown are mean of two independent experiments, standard deviation is presented as error bars. Results are shown for **ABC**. 2h protocol, **DEF**. Overnight protocol **A&D**- Growth curves with the high concentration of antibiotic (in orange in Fig. S1C&D). **B&E**- Corresponding total PG amount expressed as a percentage of the untreated control. **C&F**- Precursors quantification expressed as a percentage of the untreated control. Each precursor is presented as the sum of the non-amidated and amidated form (UDP-tri=UDP-M3+UDP-M3-NH_2_, UDP-penta=UDP-M5+UDP-M5-NH_2_).

Addition of the antibiotics to low density cultures induced a longer (dose-dependent) lag phase after which cells grew in the presence of the antibiotic and reached a similar final optical density to untreated cultures (**Fig. 2D, Fig. S3**). Tunicamycin was again the only exception: it drastically reduced both growth rate and final optical density reached (**Fig. 2D, Fig. S3F**). The total amount of PG was still more strongly reduced for all compounds with the exception of bacitracin-treated cells, which had a normal PG amount (**Fig. 2E, Fig. S4**). In *B. subtilis*, bacitracin is known to induce the expression of several resistance modules that jointly maintain CW homeostasis (Radeck et al. 2016). After 17 h of growth in the presence of the antibiotics, the amount of intracellular soluble precursors was however back to normal for most antibiotics (**Fig. 2F** and **Table S1B**), consistent with an adaptation of cells to grow with an impaired PG synthetic pathway. Only UDP-M3 and UDP-M5 remained significantly accumulated in D-cycloserine- and tunicamycin-treated cells, respectively, but to a lesser extent than after 2h treatment (compare **Fig. 2C** and **F**).

Taken together, these results indicate that the effects of short exposure to PG synthesis inhibitors mainly reflect primary target inhibition. First, cultures stop growing upon inhibition of the PG synthetic step. PG precursors upstream the blocked step accumulate and the total amount of PG is reduced. Second, cells start lysing, presumably because the balance between PG synthesis and PG hydrolysis is disrupted as suggested by the recovery observed in the supplementation with Mg^++^. However, cells ultimately adapt to grow in the presence of the antibiotics with a strongly reduced amount of PG in their walls.

### Sacculi of cells growing in the presence of PG inhibitors are less crosslinked but have more trimers

To get a better understanding of adaptation to grow in the presence of CW antibiotics, we performed a detailed analysis of muropeptide composition. Ultraperformance liquid chromatography (UPLC) analysis coupled to mass spectrometry (MS) allowed the identification of the main muropeptide species in our *B. subtilis* PG profile (**Fig S5A**). The main resolved monomeric species included disaccharide-dipeptide (M2) and -tripeptide (M3) and M3 amidated in the *m*-DAP (M3-NH_2_). Disaccharide-pentapeptide with one amidation in the *m*-DAP (M5-NH_2_) co-eluted with the M2 muropeptide. Based on MS quantification, M5-NH_2_ contributes to only 0.2% of the peak (M2: 99.8%) in untreated cells. All main dimers contained 4→3 crosslinks (resulting from DD-TP) and were of type D43 (disaccharide tetrapeptide -donor- with the terminal D-Ala linked to the carboxyl terminal *m*-DAP of a disaccharide tripeptide -acceptor-), with one or two *m*-DAP amidation(s) and de-N-acetylated or not, as previously reported (Warth et Strominger 1971; Atrih et al. 1999). The muropeptides eluting in peaks 7 and 8 have the same mass and amino acid composition (D43-NH_2_) (**Fig. S5A**). We believe these are isoforms and the difference in retention is due to the position of the amide group, on either *m*- DAP of the dimer. Additionally, the mass of peak 9 ([M+1] =1790.8) could correspond to a D43 with 4 amidations (D43-4*NH_2_; mass error of 18.9 ppm). MS fragmentation did not allow us to identify where the two putative additional amidations are, possibly on the D-Glu (yielding D-isoglutamine) as this is common in many species (Warth et Strominger 1971; Vollmer, Blanot, et De Pedro 2008). However, we did not detect any monomer (M3-NH_2_) amidated on the D-Glu, and if there was any monomer doubly amidated (M3-2*NH_2_) it would most likely have eluted too early in our chromatogram profiles to be detected. Finally, two major trimeric species of type T443 (disaccharide tetrapeptide disaccharide tetrapeptide disaccharide tripeptide), with two or three amidations (T443-2*NH_2_ and T443-3*NH_2_, respectively) were also resolved (**Fig. S5A**).

Based on the quantification of these muropeptide species for each antibiotic condition (**Fig. S5B-C** and **Table S2**), we first assessed overall structural features and covalent modifications of the PG (**Fig. 3A**). For all PG inhibitors, monomers importantly accumulated while dimers importantly decreased, in particular after overnight treatment, reducing the cross-linking of the sacculus and suggesting reduced TP and/or increased activity of EPs. Sacculi were greatly enriched in monomers resulting from both DD-EP and DL-EP activity (**Fig. 3A and Fig. 4A-B**). Another noticeable feature was accumulation of trimers after 2h treatment with PG precursors inhibitors and after overnight treatment with bacitracin and TP/TG inhibitors. Trimers are thought to have a relevant structural role as they are hubs connecting multiple glycan strands (Vollmer, Blanot, et De Pedro 2008). They have been suggested to represent the linking point of short-to-long glycan strands (Vollmer, Blanot, et De Pedro 2008), and also to help strengthening the sacculus when glycan strands are short (de Pedro et Cava 2015). Since cells grown in the presence of TP/TG inhibitors have down to 10% of PG relative to untreated cells (**Fig. 2E** and **Fig. S4B**), cross-linking of trimers may help to ensure sacculus integrity. Trimers are more compact than dimers and thus they may also be less accessible to the activity of EPs X) (de Pedro et Cava 2015).

**Figure 3.**
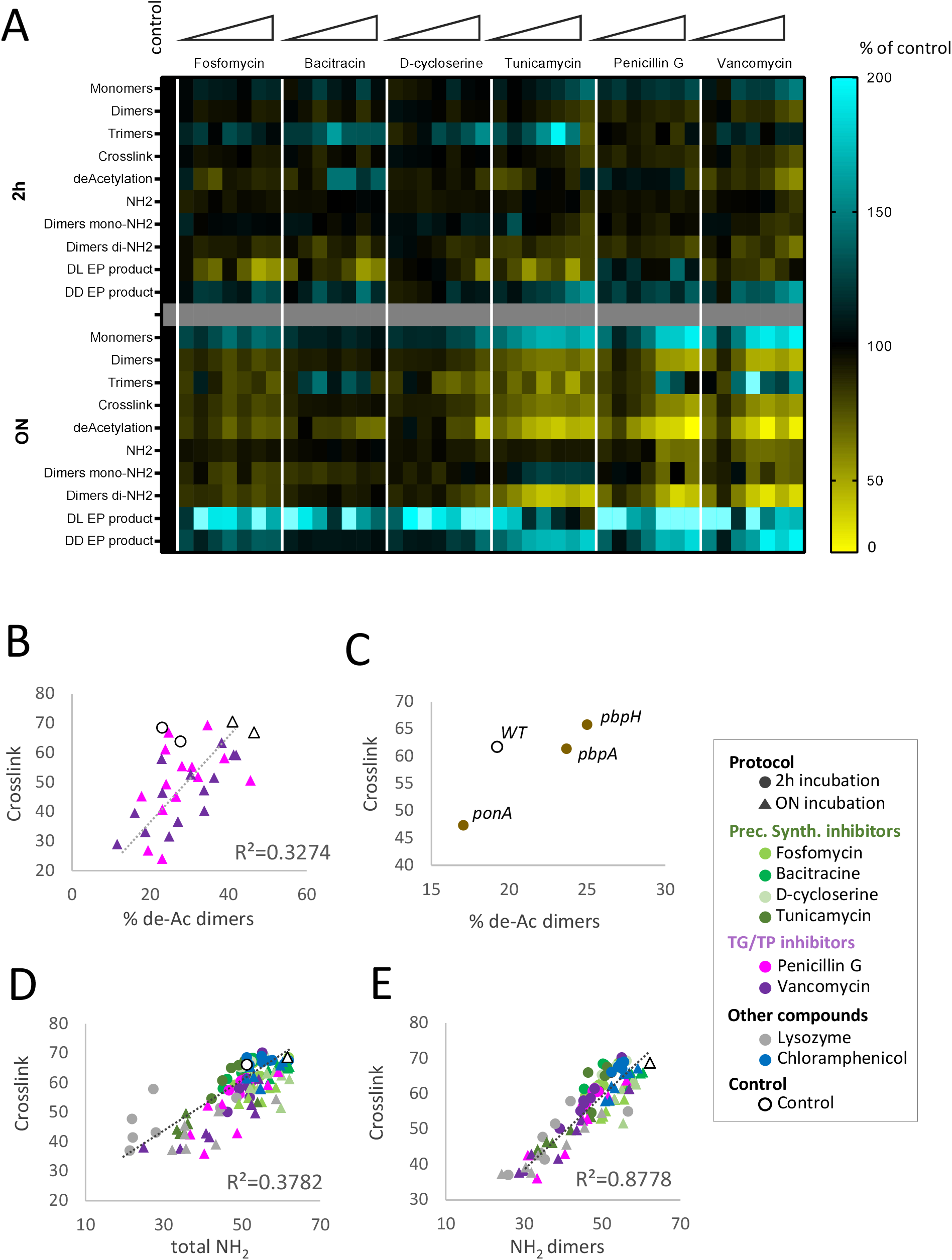
Antibiotic treatment reveals a correlation between PG crosslink, amidation and de-N-acetylation. **A.** Heatmap of PG features. Each PG feature is expressed as a percentage of the control. PG analysis was carried out for all the antibiotic’s concentrations in Fig S1C&D. Light cyan: above 220% **B.** Correlation between crosslinking (expressed as % of total PG) and de-N-acetylation (expressed as % of dimers) for TG/TP inhibitors, for the ON protocol. De-acetylation of dimers = (peak 4+ peak 5)/(peak4 + peak5 + peak6 + peak7 + peak8 + peak9) **C.** Correlation between crosslinking (expressed as % of total PG) and de-N-acetylation (expressed as % of dimers) of mutant strains. Each value is a mean of three independent experiments. De-acetylation of dimers = (peak 4+ peak 5)/(peak4 + peak5 + peak6 + peak7 + peak8 + peak9) **D.** Correlation between crosslinking (expressed as % of total PG) and amidation (expressed as % of total possible amidation) for all antibiotics, for the two protocols. Amidation = % of amidated species vs all possibly amidated species = 100* (peak1 + 0.002*peak3 + 2*peak4 + peak5 + 2*peak6 + peak7 + peak8 + 2*peak9 + 2*peak10 + 3*peak11) / ((peak1 + peak2 + 0.002*peak3) + 2*(peak4 + peak5 + peak6 + peak7 + peak8 + peak9) + 3*(peak10 + peak11)) **E.**Correlation between crosslinking (expressed as % of total PG) and amidation of dimers (expressed as % of total possible amidation of dimers) for all antibiotics, for the ON protocol. Amidation Dimers = 100* (2*peak4 + peak5 + 2*peak6 + peak7 + peak8 + 2*peak9) / (2*(peak4 + peak5 + peak6 + peak7 + peak8 + peak9))

**Figure 4.**
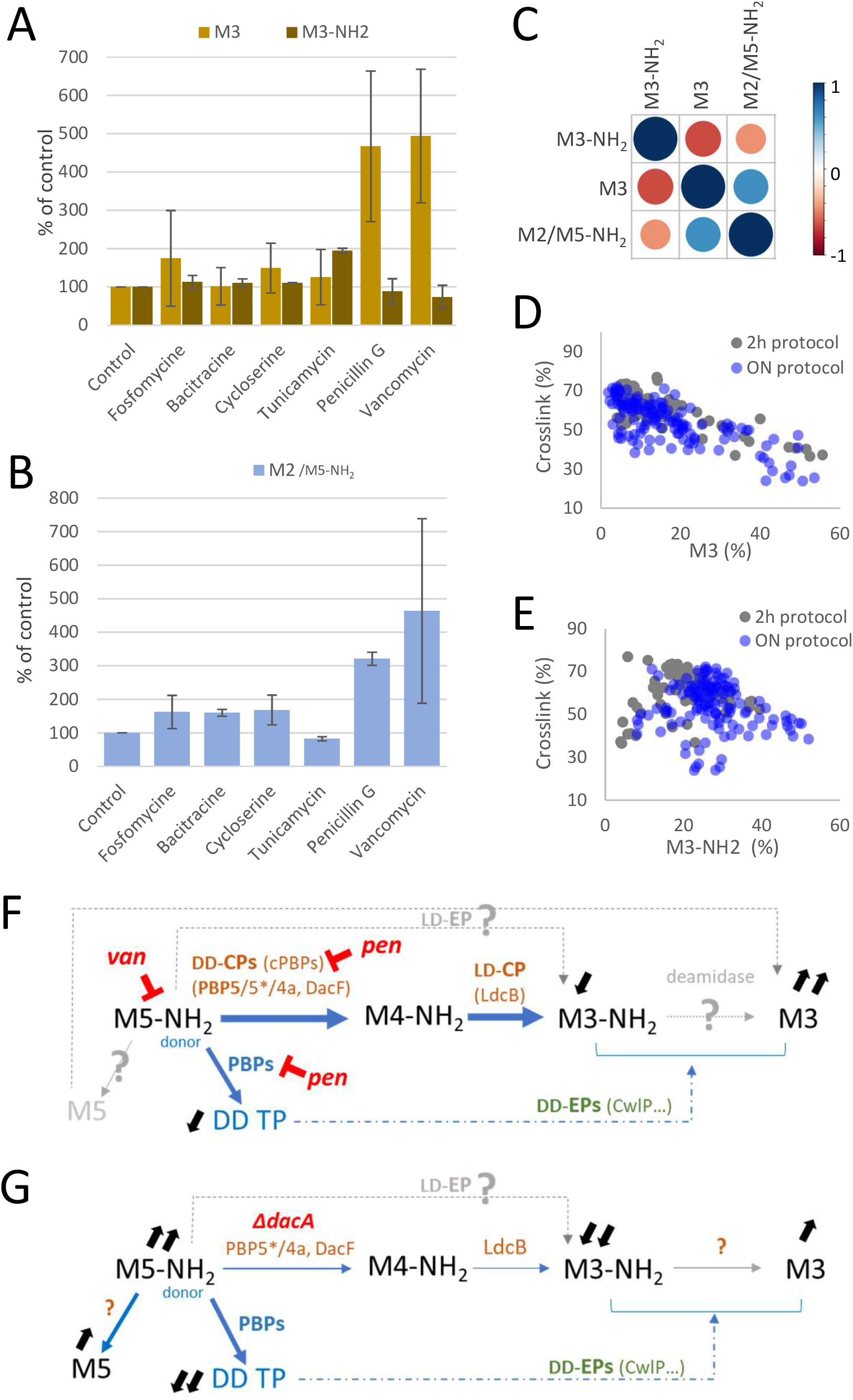
Non-amidated M3 accumulates with antibiotics inhibiting TP reaction, and is anti-correlated with amidated M3 and non-amidated M2. **A, B**. Quantification of M3 and M3-NH_2_ (A) and M2 (B) monomers for the overnight protocol, for the high concentration user in precursors analysis (in orange in Fig S1D), expressed as a percentage of total PG features. Results shown are mean of two independent experiments, standard deviation is presented as error bars. **C**. Matrice of correlation of monomers for the high concentration of antibiotic used for the PG precursors quantification (in orange in Fig S1C&D), for all antibiotics, for the 2 protocols, pooled together. **D, E**. Correlation between crosslinks and M3 monomers (D) and between crosslink and M3-NH_2_ monomers (E) expressed as a percentage of total PG features (mean of two independent experiments). **F, G**. Relations between the monomers and their respective variations, for penicillin- and vancomycin-treated cells (F) and for *dacA* mutant (G).

### Adaption to CW antibiotics involves a reduction in de-N-acetylation and amidation of the sacculus

We also assessed changes in amidation and N-acetylation, which have been shown to participate in CW homeostasis by modulating the autolytic activity of PG hydrolases (Moynihan et Clarke 2011; Yadav, Espaillat, et Cava 2018; Dajkovic et al. 2017). Note that O-acetylation was not detected in our analysis. Our PG extraction protocol involves treating samples at basic pH (8.5-9) for reduction and then adjusting to acidic pH (3-4) (Alvarez et al. 2016). Since O-acetyl groups are labile when subjected to basic or acidic pH, they were likely stripped out (Guariglia-Oropeza et Helmann 2011). Amidation of *m*-DAP and N-acetylation of GlcNAc occur intracellularly, presumably at the level of *m*-DAP of lipid II (Atrih et al. 1999; Vollmer, Blanot, et De Pedro 2008; Dajkovic et al. 2017). Newly externalised lipid II precursors are 100% amidated and N-deacetylated (Vollmer, Blanot, et De Pedro 2008; Hoyland et al. 2014) and enzymes with de-amidase and de-acetylase activity regulate these modifications in the sacculus. De-N-acetylation (de-Ac) abolishes binding of LysM domains to polysaccharides containing GlcNAc including PG (Mesnage et al. 2014). Since LysM domains are often found in PG hydrolases, it is thought that de-Ac reduces their binding to PG, and thus their activity (Rae et al. 2011; Yadav, Espaillat, et Cava 2018). *m*-DAP amidation was shown to allow physiological control over PG hydrolysis in *B. subtilis*; it was proposed that decreasing the amount of doubly amidated dimers in the sacculus may allow to increase the activity of one or more PG hydrolases (Dajkovic et al. 2017; Tesson et al. 2022).

The sacculi of untreated cells were less de-N-acetylated and less amidated during exponential growth (2h protocol) than in stationary phase (overnight protocol) (**Fig. 3BD** and **Fig. S6A**, open symbols, and **Table S2**), consistent with the key role of PG hydrolases in active CW growth. In the presence of CW antibiotics, changes in these two covalent modifications were more important after overnight treatment (**Fig. 3A**), suggesting that they may be part of the adaptative response to the presence of the antibiotics. Sacculi were less de-N-acetylated for all antibiotic treatment, while total amidation (of all muropeptide species: monomers, dimers and trimers) was strongly reduced in penicillin G and vancomycin-treated cells. Interestingly, the amount of singly amidated dimers was only slightly affected by most antibiotics, while the amount of doubly amidated dimers was significantly reduced, especially in cells grown overnight in the presence of tunicamycin, vancomycin and penicillin G (**Fig. 3A**).

When we pooled together the results for all concentrations of each antimicrobial compound, for the 2 protocols, two correlations emerged. First, crosslink was correlated with total de-Ac, mostly after overnight adaptation (**Fig. S6A-B**). This was expected since de-Ac is calculated as the relative amount of de-N-acetylated muropeptides, and this modification was only detected in dimers in our chromatograms (**Fig. S5A**). Since total crosslink was calculated as dimers+2*trimers, the two parameters are directly proportional. When we calculated instead the percentage of de-Ac within dimers (% of de-Ac dimers relative to all dimers), a correlation was still found after overnight adaptation (**Fig. S6C-D**), in particular in the presence of vancomycin and penicillin G (TP inhibitors) (**Fig. 3B** and **Fig. S6E-I**). Such correlation might suggest that cells growing in the presence of TP inhibitors reduce de-Ac to -counter-intuitively-upregulate PG hydrolases, which further reduce crosslinking. Alternatively, reduced de-Ac could be a consequence of reduced crosslinking. In this scenario, de-Ac could be functionally related to TP, with TP by PBPs preceding de-Ac of dimers. Then, if TP is inhibited de-Ac would be reduced. To test this hypothesis, we analysed the muropeptide composition of TP mutants including *ΔponA* (encoding PBP1, the major aPBP of *B. subtilis*), previously reported to have a sacculus less crosslinked than wild-type cells (Atrih et al. 1999), *ΔpbpH* and *ΔpbpA* (encoding the co-essential PbpH and PBP2A, respectively, major bPBPs). The sacculi of *ΔponA* mutant cells were less crosslinked and, as hypothesized, contained less de-N-acetylated dimers relative to the wild-type (**Fig. 3C**). The PG of the *ΔpbpH* mutant was more crosslinked, possibly because of deregulation of PBP2A, and dimers significantly more de-N-acetylated. However, dimers were also more de-N-acetylated in the *ΔpbpA* mutant but its crosslinking was similar to the wild-type (**Fig. 3C**). Whether the TP and de-N-acetylation reactions are functionally coupled will require further investigation.

We also found a significant correlation between crosslinking and amidation of the sacculus, in particular amidation of dimeric muropeptides, for all antibiotics and mostly upon overnight adaptation too (**Fig. 3D-E**). The reduction in crosslinking was correlated with a reduction in both singly and doubly amidated dimers for all antibiotics (**Fig. S7**). By de-amidating dimers, cells are thought to increase the activity of PG hydrolases (Dajkovic et al. 2017; Tesson et al. 2022). Based on these results, it is tempting to speculate that reduced crosslinking could be a consequence increased PG hydrolytic activity, and required for growth in the presence of PG synthesis inhibitors.

### Antibiotics inhibiting TP reactions induce a strong accumulation of non-amidated M3

To gain insight into muropeptides de-amidation and a possible upregulation of PG hydrolases in the presence of CW antibiotics, we analysed in detail the variations of individual muropeptide species. The heatmap in **Figure S8** further shows that long-term exposure to PG synthesis inhibitors triggers a modification in muropeptide composition more pronounced than a 2h treatment. The most striking variation was the accumulation of non-amidated M3 in most conditions (**Fig. S8** and **Table S2**). This increase was particularly important in cells treated overnight with lysozyme, vancomycin or with high doses of penicillin G, which also accumulated M2 significantly (**Fig. 4A-B**, **Fig. S8** and **Table S2**). Matrices of correlation between the different datasets showed that PG inhibitors in general produced a correlated accumulation of peak 3 (coelution of M2 and M5-NH_2_) and M3 monomers that anticorrelated with changes in the amount of M3-NH_2_ (**Fig. 4C** and **Fig. S9**). Furthermore, we found an inverse correlation between crosslink and M3 for all PG synthesis inhibitors and conditions (**Fig. 4D**), while no significant correlation was found between crosslink and M3-NH_2_ (**Fig. 4E**), indicating that M3 specifically accumulates when peptide crosslinks are reduced. This correlation was primarily given by the compounds that more strongly affected crosslink: lysozyme, penicillin G and vancomycin (**Fig. S10A**), which also brought about the strongest reduction in amount of PG in the sacculus (**Fig. 2E**). Penicillin G and vancomycin inhibit TP/DD-CP and TG/TP reactions, respectively, on the newly externalised PG precursor, uncrosslinked amidated disaccharide pentapeptide (M5-NH_2_) (**Fig. 4F**), which would be expected to accumulate in these conditions. M5-NH_2_ co-eluted with M2 in our chromatograms (99.8:0.2 M2:M5NH_2_ in untreated cells) (**Fig. S5**, peak 3) and the corresponding peak increased in penicillin G- and vancomycin-treated cells (**Fig. 4B**).

Since penicillin G blocks both DD-TPs and DD-CPs (**Fig. 4F**), the observed accumulation of M3 might result from either (a) de-amidation of M5-NH_2_ into M5 by a yet unknown de-amidase followed by hydrolysis of M5 directly into M3 by a LD-EP (CwlK), or by residual DD-CP (PBP5, PBP5*, PBP4a, DacF) (Smith, Blackman, et Foster 2000) and then LD-CP (LcdB), or (b) hydrolysis of M5-NH_2_ into M3-NH_2_ first followed by de-amidation of M3-NH_2_ into M3 by the same set of enzymes. To further investigate this, we analysed the muropeptide profile of a *ΔdacA* (**Fig. S11**). *dacA* encodes PBP5, the major DD-CPase of *B. subtilis* (Atrih et al. 1999) and in its absence cells greatly accumulate M5-NH_2_, which becomes the dominant muropeptide (**Fig. S11**) (Atrih et al. 1999). Interestingly, de-amidated M5 was detected in *ΔdacA* mutant cells (**Fig. S11** and **Table S2**), indicating that de-amidation can occur in uncrosslinked pentapeptide stems (**Fig. 4G**). However, M3 accumulated less than 2-fold in the *ΔdacA* mutant (**Fig. S11**), while in penicillin-treated and vancomycin-treated cells it accumulated up to 2.8 and 4.4-fold, respectively, in the same conditions. Thus, accumulation of M5 did not correlate with an accumulation of M3, further suggesting that M5 is not the main transient intermediate of M3. The strong anticorrelation found between M3-NH_2_ and M3 monomers (**Fig. 4C**) supports the idea that accumulation of M3 results from de-amidation of M3-NH_2_ into M3. We concluded that inhibition of TP reactions either diverts the consumption of M5-NH_2_ into M3-NH_2_ by the action of CPs or LD-EPs and M3-NH_2_ is then de-amidated into M3 or, alternatively, that accumulation of M3 in these conditions results from increased DD-EP activity on dimers. In the latter case, the M3-NH_2_ → M3 step may still be at play since only M3 accumulated while M3-NH_2_ decreased, yet both are expected products of DD-EPs.

It would be interesting to analyse PG composition of *dacA*, *ldcB* and *cwlP* mutants (**Fig. 4F,G**) in the presence of TG/TP inhibitors to conclude about the origin of M3.

## Conclusions

Our study revealed that cells adapt to the presence of CW antibiotics in an unexpected way: they reduce their PG amount, decrease the degree of de-N-acetylation and of amidation of dimers. This suggests that an increase of PG hydrolase activity is required for cell survival in presence of PG inhibitors. Finally, the decrease in PG crosslinking triggered by CW antibiotics correlates with an accumulation of M3, giving us some insight about the mechanisms at play during cell adaptation to PG inhibitors. These findings highlight a cellular response to primary target inhibition that leads to adaptation and tolerance to grow in the presence of PG inhibitors.

## Materials and Methods

### General procedures and growth conditions

*Bacillus subtilis* 168 wild-type strain was grown in LB medium at 30°C for overnight cultures and at 37°C for day cultures. Growth protocols are detailed in the results section. The Δ*ponA*, Δ*dacA*, Δ*pbpA*, Δ*pbpH*, Δ*ldcB* mutants are from the *Bacillus* Knock-Out Kanamycin library (Koo et al. 2017).

### Antibiotics

Stock solution of antibiotics were prepared as follows: fosfomycin 10 mg/mL in H_2_O, bacitracin 10 mg/mL in H_2_O, D-cycloserine 10 mg/mL in H_2_O, tunicamycin 10mg/mL in H_2_O with a few drops of 1 M NaOH, penicillin G 10 mg/mL in H_2_O, vancomycin 1 mg/mL in H_2_O, lysozyme 10 mg/mL in H_2_O, chloramphenicol 10 mg/mL in ethanol.

### Growth protocols & growth curves

Two growth protocols were used in the study **(Fig. S1A)**. For the “2h protocol” an overnight culture grown in LB at 30°C was diluted in LB to OD_D600_ 0.01 and incubated at 37°C in a plate reader with reading of optical density at 600 nm until mid-exponential phase (OD ~ 0.2, t0); drug was added at t0 and cells were incubated again in the plate reader for 2h before collection. For the “ON protocol”, a pre-culture was realized during the day in LB at 37°C and diluted to OD_D600_ 0.01 in LB; drug was added at the same time as dilution (t0) and growth was carried O.N. (~17h) at 37°C in a plate reader with reading of optical density at 600nm. Two independent replicates (cultures, extraction and quantification) were realized for each condition.

Analysis of the PG of the mutant strains was done in triplicate; the wild-type and mutants were grown in LB until exponential phase when the samples were taken.

### Intracellular soluble precursor preparation

Sample preparation of intracellular soluble PG samples was performed as follows (Hernández et al. 2020). Cells were grown according to the “2h protocol” and the “overnight protocol’ described above; OD was measured prior to sample collection for further normalization of the results. 500μL of cells were then centrifuged and pellets washed twice in 500 μL 0.9% NaCl before freezing. Pellets were resuspended in 100 μL water and autoclaved for 20 min at 120°C. After centrifugation the supernatant was recovered and filtered.

Soluble muropeptides were detected and characterized by MS/MS analysis as follows. Identity of the peaks was confirmed using a Xevo G2-XS QTof system (Waters Corporation, USA). The instrument was operated in positive ionization mode. Detection of muropeptides was performed by MS^E^ to allow for the acquisition of precursor and product ion data simultaneously, using the following parameters: capillary voltage at 3.0 kV, source temperature to 120 <u>°</u>C, desolvation temperature to 350 <u>°</u>C, sample cone voltage to 40 V, cone gas flow 100 l/h, desolvation gas flow 500 l/h and collision energy (CE): low CE: 6 eV and high CE ramp: 15-40 eV. Mass spectra were acquired at a speed of 0.25 s/scan. The scan was in a range of 100–2000 m/z. Data acquisition and processing was performed using UNIFI software package (Waters Corp.). An in-house compound library built in UNIFI was used for detection and identification of soluble muropeptides. Quantification was performed by integrating peak areas from extracted ion chromatograms (EICs) of the corresponding m/z value of each muropeptide.

Soluble precursors analysis was performed in biological duplicates.

### Muropeptide preparation

High-throughput PG isolation, muropeptide separation and quantification were realized essentially as described in (Hernandez et al. 2022). Briefly, sacculi were isolated by resuspension of 500μl cell pellet in LB with 1.6% SDS and autoclaving 15 min at 120°C and 1 atm. Boiled sacculi were transferred to 96-well-filter plates (AcroPrep Advance 96-well 0.2 μm GHP membrane filter plates, 2 ml volume, PALL) and washed 3 times with 800 μL water; covalently bound proteins were removed by adding 200 μL Tris 100mM pH8 and 0.4μL of proteinase K to the filters and incubating 30 min at 37°C. After an additional wash with 700μL water and centrifugation, the filter plate was transferred to a clean deepwell collecting plate. PG digestion was realised *in situ* with 1μL of muramidase at 1mg/mL in 50μL of water, O.N. at 37°C. Eluted muropeptides were collected in the collection plate and submitted to sample reduction. pH was adjusted to 8.5-9 by addition of 8μL borate buffer 0.5M pH9 and treated with 25μL NaBH_4_ 2M for 30min RT. Reaction was stopped and pH adjusted to 2.0-4.0 with orthophosphoric acid 25%.

Muropeptides were analyzed by UPLC, as previously described (Alvarez et al. 2020). Briefly, muropeptides were separated using an UPLC system (Waters) equipped with a trapping cartridge precolumn (SecurityGuard ULTRA Cartridge UHPLC C18 2.1 mm, Phenomenex) and an analytical column (Waters ACQUITY UPLC BEH C18, 130Å, 1.7 μm, 2.1 mm×150 mm), using as solvents 0.1% (v/v) formic acid in Milli-Q water (buffer A) and 0.1% (v/v) formic acid in acetonitrile (buffer B) in a 14 min run. Muropeptides were detected by measuring the absorbance at 204 nm. Peak identities were confirmed by MS/MS as described above.

Muropeptide analysis was performed in biological duplicates for each condition.

### Chromatographic data analysis and quantifications

Collected UPLC chromatographic data were analyzed using the chemometric-based pipeline described in (Kumar, Espaillat, et Cava 2017). Raw data were trimmed and baseline corrected. Then, data sets were aligned by segments using the COW algorithm.

The total PG amount was calculated as the sum of all peak areas, and normalized using the OD600 at the time of collection. Relative values were calculated using the untreated sample as reference.

The relative area of each muropeptide was calculated by dividing its peak-area by the total area of the chromatogram.

The different PG features were calculated as follows:

| Peak | Muropeptide | Identity |
| --- | --- | --- |
| 1 | M3-NH<sub>2</sub> | Disaccharide tripeptide with 1 amidation |
| 2 | M3 | Disaccharide tripeptide |
| 3 | M2/M5-NH<sub>2</sub> | Disaccharide dipeptide / Disaccharide pentapeptide with 1 amidation |
| 4 | D43-2*NH<sub>2</sub>,deAc | Disaccharide tripeptide disaccharide tetrapeptide with 2 amidations, missing 1 acetyl group |
| 5 | D43-NH<sub>2</sub>,deAc | Disaccharide tripeptide disaccharide tetrapeptide, missing 1 acetyl group |
| 6 | D43-2*NH<sub>2</sub> | Disaccharide tripeptide disaccharide tetrapeptide with 2 amidations |
| 7 | D43-NH<sub>2</sub> | Disaccharide tripeptide disaccharide tetrapeptide with 1 amidation |
| 8 | D43-NH<sub>2</sub> | Disaccharide tripeptide disaccharide tetrapeptide with 1 amidation |
| 9 | D43-4*NH<sub>2</sub> | Disaccharide tripeptide disaccharide tetrapeptide with 4 amidations |
| 10 | T443-2*NH<sub>2</sub> | Disaccharide tripeptide disaccharide tetrapeptide disaccharide tetrapeptide with 2 amidations |
| 11 | T443-3*NH<sub>2</sub> | Disaccharide tripeptide disaccharide tetrapeptide disaccharide tetrapeptide with 3 amidations |

Monomers = peaks 1+2+3

Dimers = peaks 4+5+6+7+8+9

Trimers = peaks 10+11

Crosslink = (peak 4+ peak 5+ peak 6+ peak 7+ peak 8+ peak 9)+2*(peak 10+ peak 11)

De-acetylation = peaks 4+5

De-acetylation of dimers = (peak 4+ peak 5)/(peak4 + peak5 + peak6 + peak7 + peak8 + peak9) DL-EP products = M2 = peak 3

DD-EP products = M3 + M3-NH2 = peaks 1+2

Total EP products = peaks 1+2+3

Amidation = % of amidated species vs all possibly amidated species = 100* (peak1 + 0.002*peak3 + 2*peak4 + peak5 + 2*peak6 + peak7 + peak8 + 2*peak9 + 2*peak10 + 3*peak11) / ((peak1 + peak2 + 0.002*peak3) + 2*(peak4 + peak5 + peak6 + peak7 + peak8 + peak9) + 3*(peak10 + peak11)) Amidation Dimers = 100* (2*peak4 + peak5 + 2*peak6 + peak7 + peak8 + 2*peak9) / (2*(peak4 + peak5 + peak6 + peak7 + peak8 + peak9)) Doubly amidated dimers / all dimers =100* (peak4 + peak6 + peak9) / (peak4 + peak5 + peak6 + peak7 + peak8 + peak9) Singly amidated dimers/ all dimers = 100*(peak5 + peak7 + peak8) / (peak4 + peak5 + peak6 + peak7 + peak8 + peak9)

### Data analysis and statistical methods

At least two independent experiments of each condition were considered and used to calculate mean and standard deviation (error bars), except for the scatterplot showing correlations **(Fig. 3B, Fig. S6, S7, Fig. 4D,E)** where all replicates are plotted. On these scatterplots, R^2^ of correlation was calculated using linear regression. Heatmaps **(Fig. 3A, Fig. S8)** were realized using GraphPad software.

### Correlation matrices

Correlations were calculated using R (version 4.1.1) in the RStudio environment -2012.09.0 / Build 351) and the correlation matrices were visualised using the ‘corrplot’ function (Wei et Simko 2021). Blue and red colours in correlation matrices indicate positive and negative correlation, respectively. The shade of the colour and the circle diameter refers to the amplitude of the correlation.

## Acknowledgments

This project has received funding from the European Research Council (ERC) under the Horizon 2020 research and innovation program (grant agreement No 772178 to R.C.-L.). Research in the Cava lab is supported by the Swedish Research Council (VR), The Knut and Alice Wallenberg Foundation (KAW), The Laboratory of Molecular Infection Medicine Sweden (MIMS) and The Kempe Foundation.

